# Sparse synaptic connectivity is required for decorrelation and pattern separation in feedforward networks

**DOI:** 10.1101/108431

**Authors:** N. Alex Cayco-Gajic, Claudia Clopath, R. Angus Silver

## Abstract

Pattern separation is a fundamental function of the brain. Divergent feedforward networks separate overlapping activity patterns by mapping them onto larger numbers of neurons, aiding learning in downstream circuits. However, the relationship between the synaptic connectivity within these circuits and their ability to separate patterns is poorly understood. To investigate this we built simplified and biologically detailed models of the cerebellar input layer and systematically varied the spatial correlation of their inputs and their synaptic connectivity. Performance was quantified by the learning speed of a classifier trained on either the mossy fiber input or granule cell output patterns. Our results establish that the extent of synaptic connectivity governs the pattern separation performance of feedforward networks by counteracting the beneficial effects of expanding coding space and threshold-mediated decorrelation. The sparse synaptic connectivity in the cerebellar input layer provides an optimal solution to this trade-off, enabling efficient pattern separation and faster learning.

---

The ability to distinguish similar, yet distinct patterns of sensory inputs is a core feature of the nervous system. Pattern separation underlies such everyday activity as recognizing faces and distinguishing odors. Early theoretical work showed that divergent excitatory feedforward networks can separate patterns of neuronal activity by projecting them onto a larger population of neurons and by reducing the fraction of neurons active, forming a ‘sparse code’ where the likelihood of overlapping patterns is low^1–6^. Divergent feedforward networks, thought to be involved in pattern separation, are widespread in the nervous system of both vertebrates and invertebrates, including the olfactory bulb^7,8^, mushroom body^9,10^, dorsal cochlear nucleus^11^ and hippocampus^12,13^. But perhaps the most well studied example is the input layer of the cerebellar cortex, which combines many different types of sensory modalities and motor command signals^14^.

The input layer of the cerebellar cortex has an evolutionarily conserved network structure, in which granule cells receive 2-7 synaptic inputs, with the claw-like ending of each of their dendrites innervating a different mossy fibre^14^. Interestingly, other divergent feedforward networks also have relatively few synapses: granule cells in the dorsal cochlear nucleus have 2-3 dendrites^15^ while Kenyon cells in the fly olfactory system have around 7 synaptic inputs^16^. This raises the question of why the synaptic connectivity of these networks is so similar. Our recent work has shown that having few synaptic inputs per granule cell provides an optimal trade-off between sparsening population activity and efficient information transmission^3^. However, there are several other important determinants of pattern separation, including “expansion recoding” (i.e., representing information in a higher-dimensional space^1,17–19^), and reducing correlations in the input patterns^9,10^. But our limited understanding of the relative importance of these properties makes the relationship between pattern separation and network structure unclear.

We examined the relationship between network structure and pattern separation in the cerebellar input layer by studying how divergent feedforward networks transform highly overlapping mossy fiber activation patterns. Using a combination of simplified and biologically detailed models, we disentangled the effects of correlations from expansion and sparsening of spatially clustered input patterns. Moreover, we quantified pattern separation performance by assaying learning speed using a machine-learning algorithm. Our results establish that the excitatory synaptic connectivity of feedforward networks is a major determinate of pattern separation performance. Furthermore, they suggest that the evolutionarily conserved sparse synaptic connectivity found in divergent feedforward networks is essential for separating spatially correlated input patterns.

## Results

The cerebellar input layer consists of mossy fibers (MFs), which form large en passant mossy-type synapses called rosettes, granule cells (GCs) which have ~4 short dendrites, and inhibitory Golgi cells which form an extensive dense axonal arbor that spans the local region. To capture the excitatory synaptic connectivity we used an anatomically accurate 3D model of a local region of the cerebellar input layer network^3^. The 80 μm diameter model had experimentally measured densities of MF rosettes (~180 in total) and GCs (~480 in total) and random connectivity, subject to the spatial constraint that MF-GC distances were close to 15 μm (Fig. 1a). Importantly, this anatomically detailed model reproduced the measured 1:2.9 local expansion ratio between MF rosettes and GCs, the 1:12 divergence at the rosette-GC synapse and the sampling of 4 different rosettes by individual GCs.

**Figure 1.**
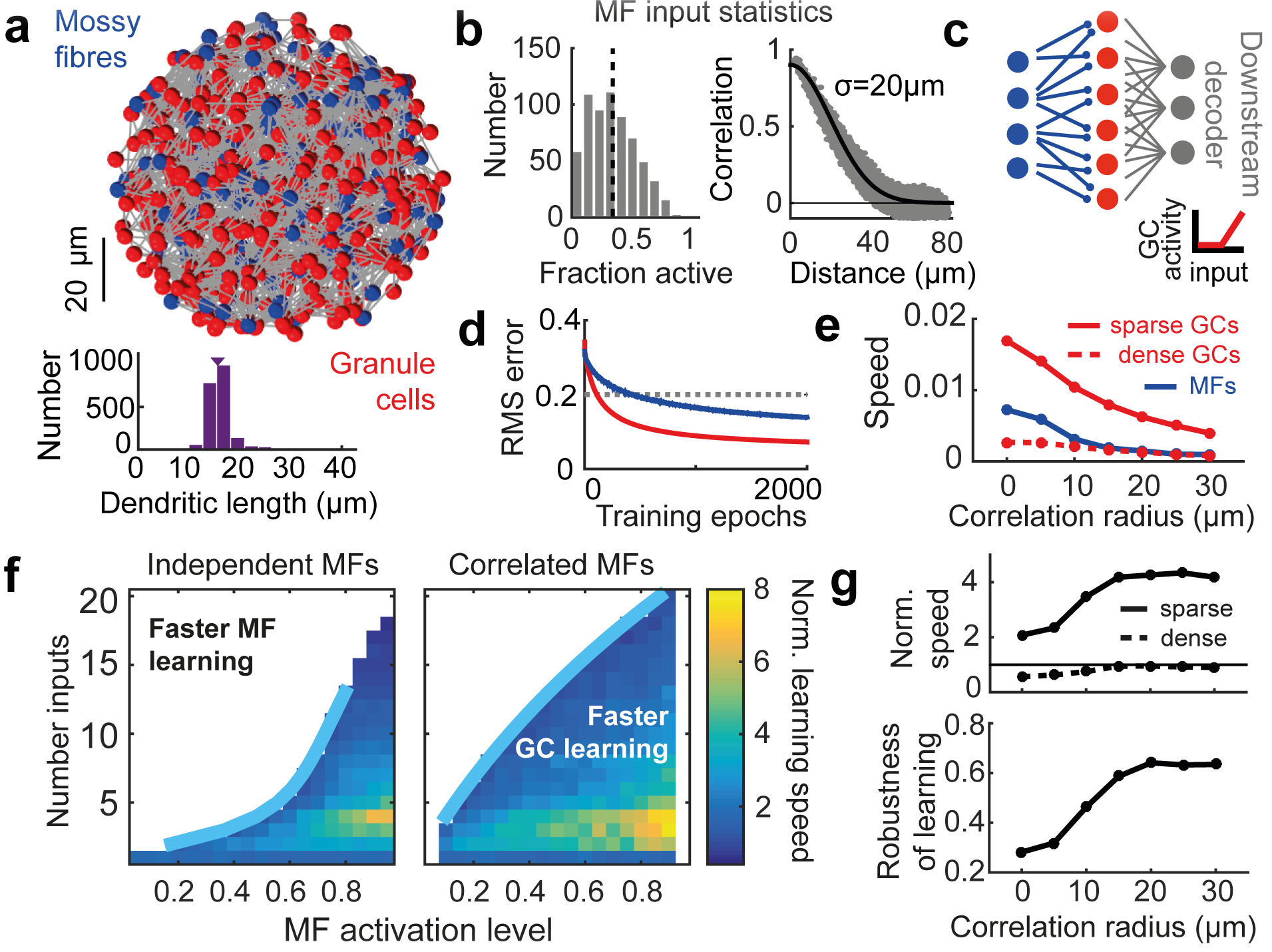
Presence of a simple feedforward model of the cerebellar input layer with sparse, but not dense, synaptic connectivity speeds learning. (**a**) Top: Anatomically constrained 3D model of cerebellar input layer. Positions of Granule Cells (GCs, red) and Mossy Fibers (MFs, blue) within an 80 μm ball. Synaptic connections are shown in gray. Bottom: Distribution of dendritic lengths. Arrow indicates mean. (**b**) Example of MF statistics generated with correlation radius of 20 μm and 30% activated MFs. Left: Histogram of the fraction of active MFs over different activity patterns. Right: Correlation between MF pairs as a function of distance between them (grey). Black lines indicated the specified average activation level (left) and spatial correlations (right). (**c**) Schematic of feedforward network model (blue MFs; red, GCs). The downstream perceptron-based decoder classifies either GC patterns (as shown) or else raw MF patterns without the MF-GC layer. Inset shows the rectified-linear GC transfer function. (**d**) Example of root-mean-square error as a function of the number of training epochs during learning based on MF (blue) or GC (red) activity patterns. Dashed line indicates threshold error. For this example, N_syn_ = 4 and 50% of MFs are activated. (**e**) Raw learning speed of perceptron classifier for different correlation radii, for MFs (blue) or GCs with sparse (solid red line, N_syn_ = 4) or dense (dashed red line, N_syn_ = 16) connectivity. (**f**) Normalized learning speed (GC speed/MF speed) shown for different synaptic connectivities and MF activation levels. Blue lines represent double exponential fit of the boundary at which the normalized speed equals 1 (i.e., when the perceptron learning speed is the same for GC and MF activation patterns). For clarity, only the region in which the normalized speed > 1 is shown. Left: independent MF activation patterns. Right: Correlated MF inputs with a correlation radius of σ = 20 μm. (**g**) Top: Median normalized learning speed (over different MF activation levels) for sparse (solid line, N_syn_ = 4) and dense (dashed line, N_syn_ = 16) synaptic connectivities, plotted against correlation radius. Bottom: Robustness of rapid GC learning for different correlation radii.

To capture spatial correlations in the MF activation patterns conveying sensorimotor information to the cerebellar cortex, we used a technique to create spike trains with specified firing rates and spike correlations^20^. A Gaussian correlation function was used to describe the distance-dependence of rosette co-activation, which was parameterized by its standard deviation σ (Fig. 1b). To explore how synaptic connectivity and input correlations affect pattern separation we varied the number of synaptic connections per GC (N_syn_) in the model and presented the networks with different activity patterns with different values of σ. We first implemented a simplified, tractable rectified-linear model of GCs with a high threshold and assayed network performance by training a perceptron decoder to classify either MF or GC population activity patterns into randomly assigned classes (Fig. 1c).

## Sparse connectivity speeds learning and increases robustness

We first tested whether the evolutionarily conserved connectivity in the cerebellar input layer (N_syn_ = 4) could separate MF activation patterns and thus aid learning. Network performance was assayed with learning ‘speed’ (quantified by the number of training epochs required to reach a threshold error level) of a downstream perceptron decoder^17^. Comparison of learning speed when the perceptron was connected to the raw MF input (blue) or the processed GC output (red) confirmed that the GC layer network speeds learning (Fig. 1d). However network performance depended strongly on input correlations and the density of connectivity (Fig. 1e). Indeed, the speed of learning for more densely connected networks (N_syn_ = 16, dashed red line in Fig. 1e) was worse than raw MF input.

To quantify the relationship between synaptic connectivity and learning speed we generated a family of network models with different N_syn_ (ranging from 1 to 20) and determined their performance across the full range of MF input activity level (i.e. the fraction of active MFs). To compare network performance across different conditions we normalized the learning speed of the classifier when connected to the GCs by the learning speed when connected directly to the MFs, such that a normalized speed >1 indicated that the GC layer improved learning performance. For independent MF activation patterns (σ = 0 μm) the normalized learning speed was substantially increased across a wide range of input activity in networks with few synaptic connections per GC (Fig. 1f, left). The normalized performance improved for higher levels of MF activation because the MF activity was too dense for efficient learning. Interestingly, the fastest speed up occurred with ~4 synapses per GC. However, as N_syn_ increased, the range of MF activity over which the GC layer network sped learning decreased.

When spatial correlations were introduced in the MF input, the ranges of MF activation and synaptic connectivity over which the GC layer improved learning speed increased. However, optimal performance (up to an 8-fold speed up) occurred when synaptic connectivity was sparse (i.e. the number of synaptic connections per GC was low; N_syn_ = 2-5; Fig. 1f, right), as for the case with spatially independent input. Normalized learning speed increased with the MF correlation radius σ but saturated at around 15 μm (Fig. 1g, top). Moreover, the fraction of the parameter space in which GC learning outperformed MF learning (a measure we refer to as ‘Robustness’ of GC learning; see Supplementary Methods) also saturated around 15 μm (Fig. 1g, bottom). These results suggest that to improve learning performance in downstream classifiers, cerebellar-like feedforward networks require sparse synaptic connectivity.

## Sparsening and expansion in coding space

To better understand why sparsely connected feedforward networks improve learning, while densely connected networks do not, we analyzed how these networks transform activity patterns. Marr-Albus theory posits two factors that are necessary for pattern separation in cerebellar cortex: sparsening and expansion recoding. We first tested whether sparse coding could explain the dependence of learning speed on network connectivity by measuring the population (i.e., spatial) sparseness of GC and MF activation patterns^21^. To compare the change in sparseness across parameters we normalized the sparseness in the GC population by the sparseness of the MF population. Although networks with many synaptic connections result in the sparsest activity^3^, the normalized sparseness was only weakly dependent on different numbers of synaptic inputs per GC (Fig. 2a, Fig. 2b, top). Moreover, increasing the correlation radius decreased the normalized sparseness and had no effect on the robustness of sparsening (cf. Fig. 2b and Fig. 1g). Therefore the change in population sparseness was not able to account for the effect of network connectivity and MF correlations on learning speed in our threshold-linear models.

**Figure 2.**
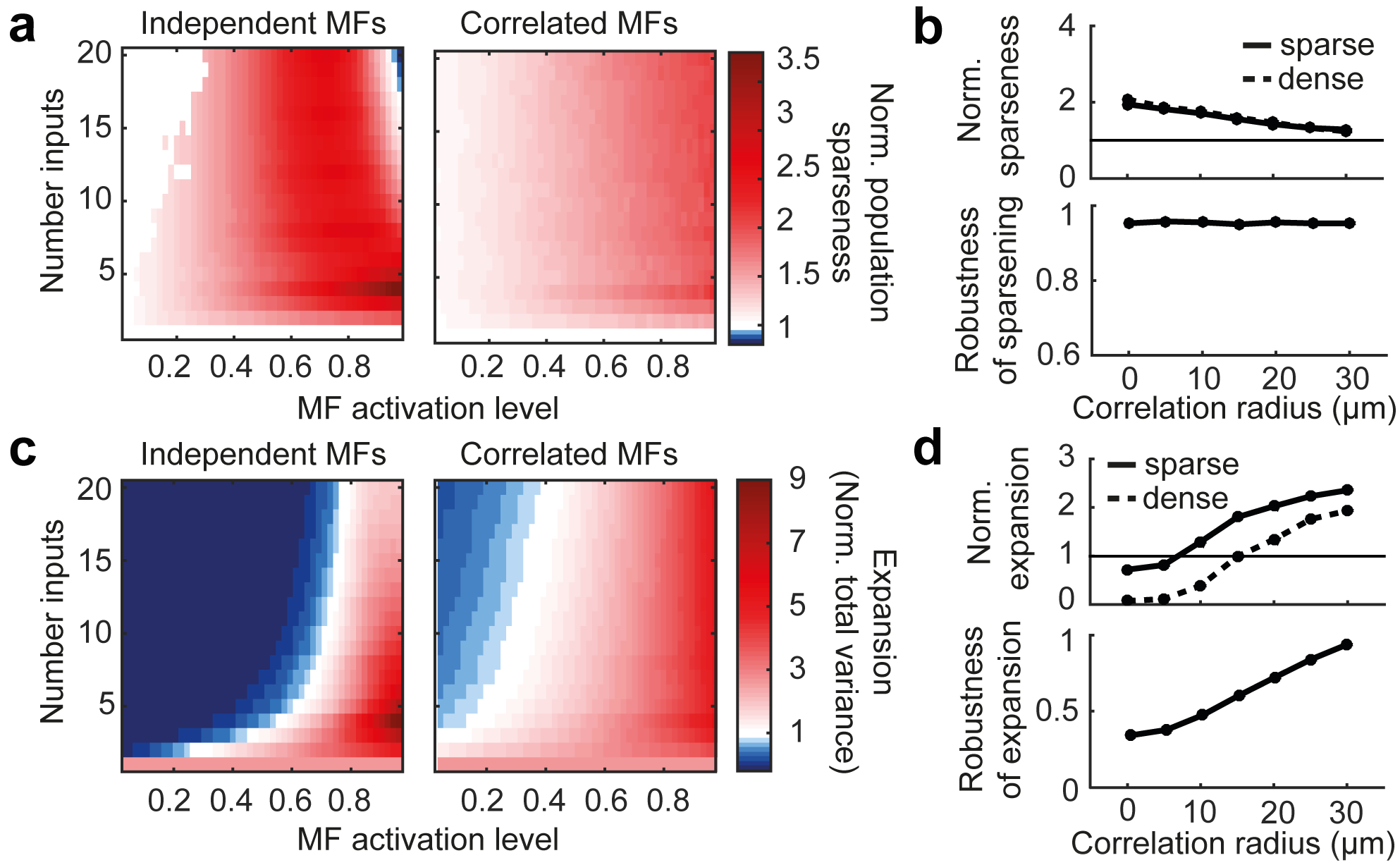
Cerebellar input layer sparsens and expands input patterns. (**a**) Normalized population sparseness (granule cell sparseness/ mossy fiber sparseness) for independent mossy fiber (MF) activation patterns (left) and correlated MF inputs (right, σ = 20 μm). (**b**) Top: Median normalized sparseness for sparse (solid line, N_syn_ = 4) and dense (dashed line, N_syn_ = 16) synaptic connectivities, plotted against correlation radius. Bottom: Robustness of sparsening for different correlation radii. (**c-d**) Same as A-B plotted for normalized total variance.

We next considered whether expansion in coding space could explain the trends in pattern separation that we have observed. Expansion recoding is thought to speed learning by increasing the distance between patterns in coding space. A key property of such expansion is the size of the distribution of activity patterns, which can be quantified by calculating the total variance in activity across the GC population (i.e., the sum of the variances), normalized by the total variance of the MF population. The normalized total variance captures both the expansion in dimensionality (i.e., due to the 1:2.9 expansion ratio) and any change in the overall size of the population coding space. The change in total variance was a better predictor of the change in learning speed than the population sparseness alone, as it was increased in the regions of parameter space where the expansion in coding space improved learning (left panels of Fig. 2c and Fig. 1f). However, the total variance could not fully explain how the network structure affected learning (assuming a linear relationship with the learning speed), since it tended to underestimate network performance for sparsely connected networks and overestimate performance for densely connected ones, particularly for correlated MF inputs (right panels of Fig. 2c and Fig. 1f). Moreover, the magnitude and robustness of the normalized total variance increased approximately linearly with MF correlations (Fig. 2d), unlike the saturation observed for learning speed (Fig. 1g). Thus the dependence of pattern separation and learning speed on network connectivity and MF input properties cannot be accounted for by changes in the sparseness and the size of the coding space alone.

## Decorrelation of MF activation patterns

The presence of spatial correlations in MF inputs is expected to reduce the dimensionality of activity patterns and slow learning due to increased pattern overlap. Mathematically, the shape of the distribution of activity patterns can be related to the covariance matrix, since the square roots of its eigenvalues correspond to the lengths of the principal directions of activity space (as illustrated for 3 dimensions in Fig. 3a, top). Independent MF activation results in more uniform eigenvalues (e.g. a sphere in 3 dimensions), whereas more correlated distributions have a more heterogeneous spread of eigenvalues and hence an elongated distribution (Fig. 3a, bottom).

**Figure 3.**
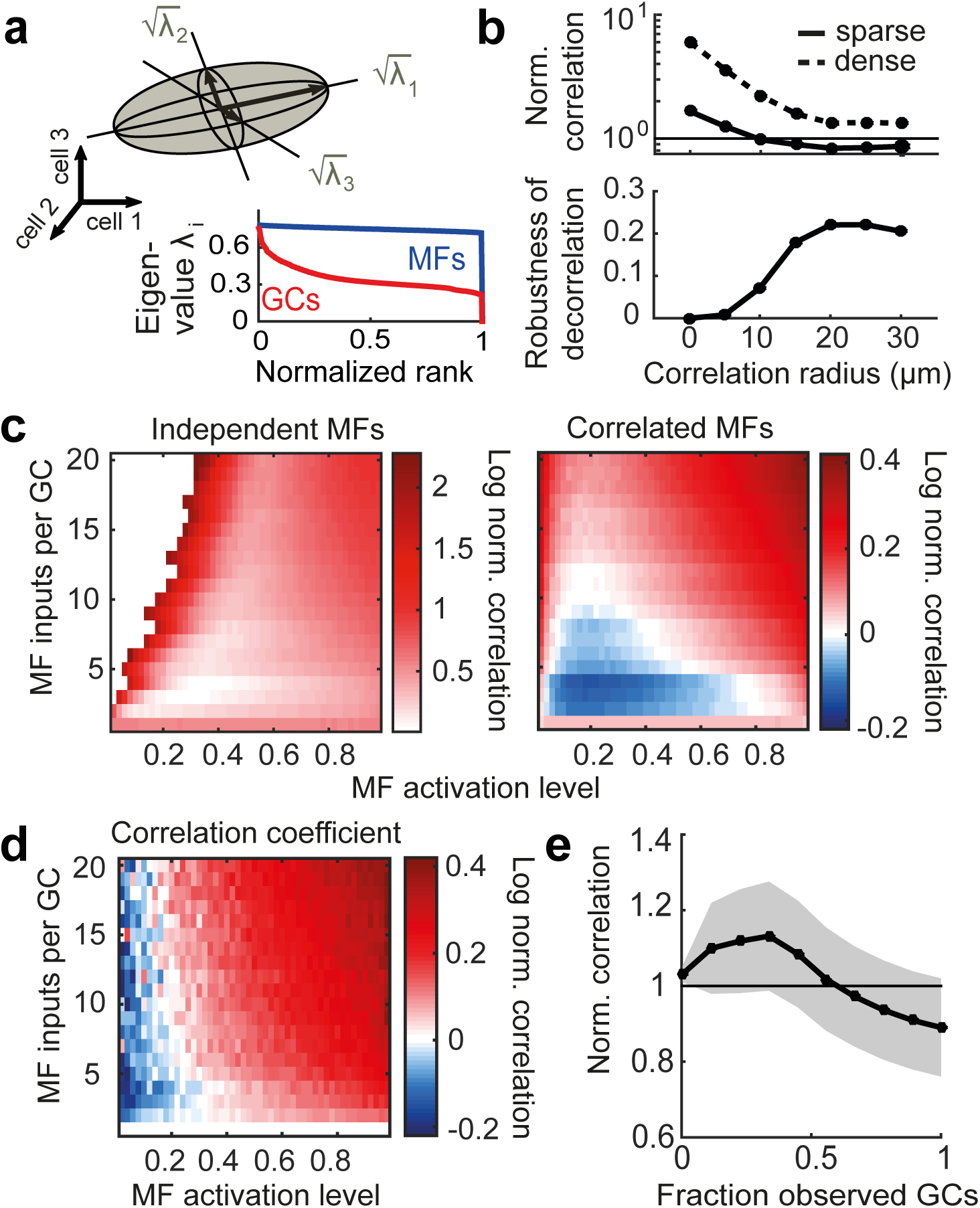
Correlations in activity increase with the extent of excitatory synaptic connectivity in feedforward networks. (**a**) Top: Illustration depicting a distribution of neural activity patterns (grey ellipsoid) in activity space. Mathematically, principal lengths (black arrows) are equal to the square roots of the eigenvalues of the covariance matrix. Bottom: Example of ranked eigenvalues for mossy fiber (MF, blue) and granule cell (GC, red) activity patterns. Rank is normalized by dimensionality. Note that the MF eigenvalues are far more uniform than the GC eigenvalues, indicating that the MF patterns are less correlated. In this example, parameters are: N_syn_ = 4, 50% activated MFs, σ = 0μm. (**b**) Top: Median normalized population correlation (GC correlation/MF correlation) for sparse (solid line, N_syn_ = 4) and dense (dashed line, N_syn_ = 16) synaptic connectivity plotted against correlation radius. Note the logscale for the population correlation. Bottom: Robustness of GC decorrelation for different correlation radii. (**c**) Log of the normalized population correlation for independent MF activation patterns (left) and correlated MF inputs (right, σ = 20 μm). Blue region in the right panel indicates region of active decorrelation of MF patterns. (**d**) Log of the normalized Pearson correlation coefficient for correlated inputs (σ = 20 μm). (**e**) Average normalized population correlation for subpopulations of increasing size. Grey error snake indicates the standard deviation across different samples and observations. For this example, N_syn_ = 4 and σ = 20 μm.

To assay neural co-variability we introduced a population-based measure of correlation, calculated using the eigenvalues of the covariance matrix, which captured the elongation of the distribution of activity patterns (see Methods). This “population correlation” varied from 0 for an uncorrelated Gaussian with identical variances (see Supplementary Methods for a discussion on heterogeneous variances) to 1 (e.g., if all neurons have identical activity). Networks with dense synaptic connectivity exhibited considerably higher normalized population correlation (i.e. GC population correlation/MF population correlation) than networks with sparse synaptic connectivity irrespective of the correlation radius (Fig. 3b). This occurred because networks with higher synaptic connectivity receive a larger number of shared inputs from MF rosettes. In the limit of full connectivity, in which each GC is connected to all MF rosettes, all GC patterns will be identical, rendering learning of different patterns impossible. Thus sparse synaptic connectivity minimizes unwanted correlations being introduced by the network structure.

Network structure was not the only factor governing the GC population correlation. Surprisingly, when the activity patterns of MF inputs were spatially correlated, the population correlation of GCs in sparsely connected networks was often *lower* than that of the MF inputs, as revealed by plotting the normalized population correlation (Fig. 3c, right). Such decorrelation of input patterns has been shown to arise from spike thresholding, which attenuates subthreshold correlations^22^. The robustness of pattern *decorrelation* in our networks saturated when the correlation radius of the activity patterns reached ~15 μm, potentially explaining the saturation in learning observed previously (Fig. 3b, bottom, cf. Fig. 1g). Moreover, varying the expansion ratio (Supplementary Fig. 1) and including adaptive thresholding to model feedforward inhibition (Supplementary Fig. 2) produced qualitatively similar results. These results suggest that correlations arising from MF input patterns, spike thresholding, and network structure all play a key role in pattern separation.

Interestingly, when network correlations were assayed with the average Pearson correlation coefficient, rather than normalized population correlation, the ability of sparsely connected networks to perform decorrelation was no longer visible (Fig. 3d). This reveals a fundamental property of the decorrelation performed by sparsely connected feedforward networks: the population correlation takes into account the shape of the distribution at the full population-level, while the correlation coefficient only considers the marginal distributions of pairs of cells, missing how they may work together to shape the full distribution. This has important implications for measuring coordinated activity in these networks, as a large fraction of cells are required to observe population decorrelation (e.g. >60% of the population for strong input correlations; Fig. 3e). Simultaneous recordings from a substantial proportion of MFs and GCs will therefore be crucial to measure the extent of decorrelation in the input layer of the cerebellar cortex.

## Determinants of expansion and decorrelation

To understand how synaptic connectivity and spike thresholding separately contribute to pattern separation, we next analyzed pattern expansion and correlation in networks of GCs with linear transfer functions (i.e. zero threshold), since under these conditions the changes in total variance and population correlation arise solely from the network structure. The total variance of linear GCs was larger than that of the MFs over the full range of parameters; however, as the number of synaptic inputs increased, the normalized total variance decreased (Fig. 4a). This decrease occurs because as N_syn_ increases, GCs average the signals across more MFs, leading to a reduction in the size of the coding space. Comparison of these results with those from networks with GCs with nonlinear spike thresholds (Fig. 2c, left) shows that the spike thresholding nonlinearity reduces both the magnitude of the coding space and its robustness (Supplementary Fig. 3). Thus, expansion in coding space is maximal for linear networks with N_syn_ = 1, but this is reduced by increasing network connectivity and by GC thresholding.

**Figure 4.**
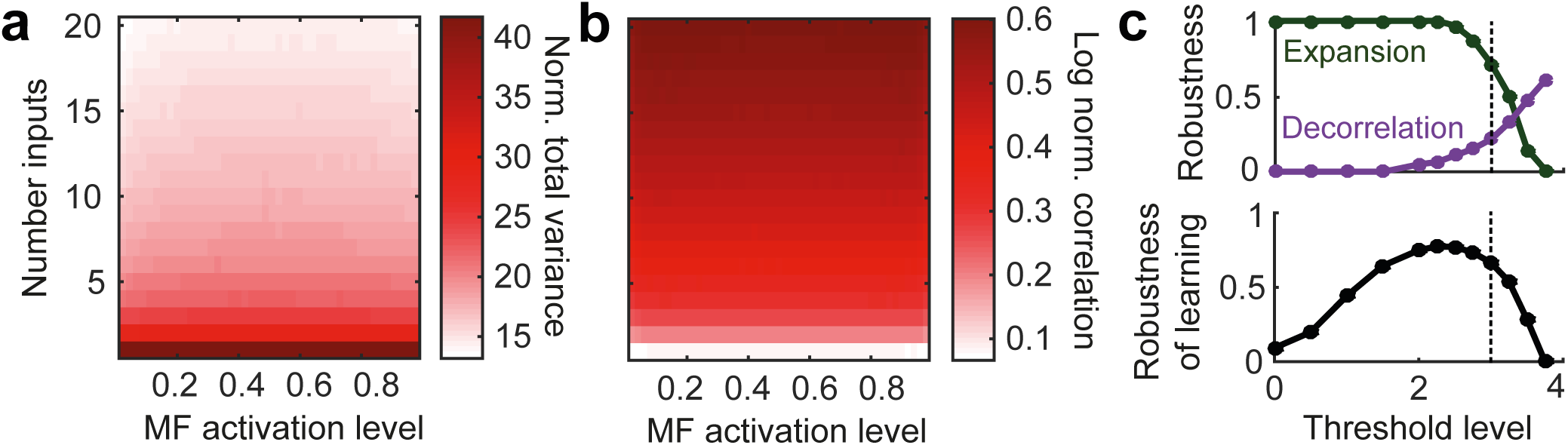
Dependence of coding space and correlation on connectivity and the role of thresholding in controlling the expansion and decorrelation. (**a**) Normalized total variance and (**b**) log normalized population correlation for networks of linear granule cells (i.e. those with zero threshold). Correlation radius is σ = 20 μm. (**c**) Top: robustness of expansion (green) and decorrelation (purple) for varying levels of granule cell (GC) threshold. Dotted line indicates the experimentally estimated value of threshold (3 of the 4 mossy fibers, MFs). Bottom: Robustness of learning for varying GC threshold.

Linear GC networks also revealed that the network structure introduces considerable population correlation (Fig. 4b). However, this was markedly reduced in networks of nonlinear neurons due to threshold-induced decorrelation (Fig. 3c). Previous work has shown that input correlations can be quenched by the presence of intrinsic nonlinearities^22^. Our results show that for feedforward networks, threshold-induced decorrelation of MF input patterns was most pronounced in sparsely connected networks (e.g. N_syn_ = 2-9). Indeed, increasing the spike threshold increased the region of decorrelation in our networks (Fig. 4c, top; see also Supplementary Fig. 3), consistent with previous work showing that sparsening of activity patterns decorrelates inputs^21,23^. In contrast, the decorrelating effect of thresholding had little effect when N_syn_ was large, due to the presence of large network-induced correlations in the inputs. Moreover, decorrelation was not observed when N_syn_ = 1, when the GC transfer function is equivalent to a linear system. Thus GC thresholding enables decorrelation of spatially correlated input patterns only when the synaptic connectivity of the network is sparse and N_syn_>1.

These simulations reveal a trade-off between expansion of coding space and a reduction of input correlations that depends on both the network connectivity and spike thresholding. Networks with dense connectivity perform pattern separation poorly because they quench coding space and introduce strong spatial correlations. By contrast, the sparse synaptic connectivity found in many feedforward networks, including the cerebellar input layer, minimizes the correlations introduced by the network, thereby enabling both expansion of coding space and decorrelation of input patterns by spike thresholding. Moreover, sparsening GC population activity by increasing spike threshold alters the trade-off between decorrelation and expansion (Fig. 4c, top). This suggests that extremely sparse codes are not optimal for pattern separation and learning due to the quenching of coding space (Fig. 4c, bottom).

## Quantifying the contributions of correlation and decorrelation

To quantify the contribution of spatial correlations to pattern separation, it was necessary to isolate their effect on learning speed from those arising from sparsening and expansion of coding space. This required ‘clamping’ the population correlation of the GC population to the value of the population correlation of the MF input. However, this constraint necessitated the removal or addition of correlations, while keeping the single-cell statistics (i.e., firing rates and variances) unchanged. To achieve this we extended methods that use random “shuffling” of the timing of activity patterns to remove all correlations^24^ by developing an algorithm that shuffled activity patterns to a pre-specified (but nonzero) level of population correlation (see Methods). The shuffled GC activity distributions had the same population correlation as the MFs (Fig. 5a), while the total variance and firing rates remained unchanged (Fig. 5b). Importantly, this procedure also maintained the sparseness of the GC population (Fig. 5c), allowing us to separate the effect of correlations from expansion and sparsening.

**Figure 5.**
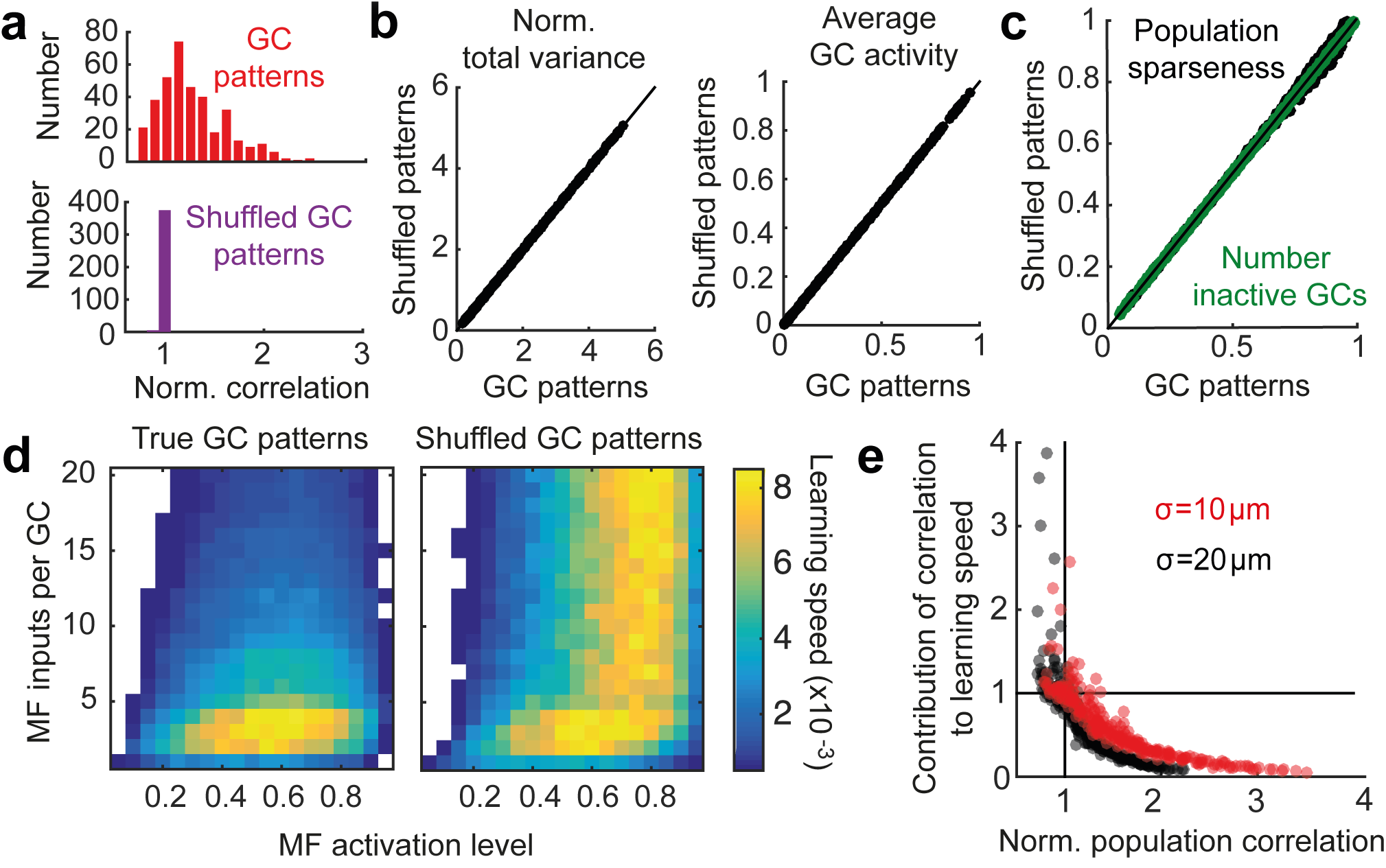
Separation of the effects of correlation on learning speed from expansion and sparsening. (**a**) Histograms of the normalized population correlation(granule cell correlation/ mossy fiber correlation) for granule cell (GC) patterns (top, red) and shuffled GC pattern (bottom, purple). The narrow distribution around 1 indicates that the shuffled GC patterns have the same population correlation as the mossy fiber (MF) patterns. (**b**) Normalized total variance (left) and average activity (right) for GC patterns (abscissa) versus shuffled GC patterns (ordinate). (**c**) Two measures of sparseness plotted for GC patterns (abscissa) and shuffled GC patterns (ordinate). Green indicates the fraction of inactive GCs; black indicates population sparseness. (**d**) Raw learning speed for true GC patterns (left) and shuffled GC patterns (right). In both panels, MF inputs are correlated with a correlation radius of σ = 20 μm. (**e**) Change in learning speed due to correlations (i.e., GC speed/shuffled GC speed) plotted against the normalized population correlation. Each point represents different values of N_syn_ and MF activation. Correlation radii were σ = 10 μm (red) or 20 μm (black).

Shuffling GC activity patterns to match population correlation in the MF input had a strong influence on learning speed when compared to the unshuffled control networks, especially for dense synaptic connectivity (Fig. 5d). Unlike the true GC responses, shuffled patterns maintained rapid learning across the full range of synaptic connectivity examined. These results confirm that correlations induced by network connectivity counteract the positive effects of pattern expansion, sparsening, and decorrelation on pattern separation and learning.

We next normalized the GC learning speed by the learning speed using shuffled GC spike patterns. This enabled us to quantify the effect correlations have on network performance after controlling for the effects of expansion and sparsening. There was a strong negative correlation between the normalized population correlation and learning (Fig. 5e), showing that population correlation reduces the normalized learning speed to as low as 0.05 (corresponding to a 20-fold reduction of learning speed). In contrast, learning speed was enhanced (up to a 4-fold increase) in sparsely connected networks where the relatively weak network-dependent correlations were quenched by spike-threshold mediated decorrelation. Thus in networks with sparse synaptic connectivity both the expansion in coding space and active decorrelation combined for faster, more robust pattern separation and learning.

## Pattern separation performed by a biologically detailed spiking model

To test the validity of the predictions from our simplified analytical models, we performed simulations with biologically detailed spiking network models of the cerebellar input layer (Fig. 6a). Active MFs were modeled as rate coded Poisson spike trains as observed in vivo^25–27^ and GC integration was modeled as an integrate-and-fire neuron with experimentally determined input resistance and capacitance, as well as AMPA and NMDA receptor-type excitatory synaptic conductances that included spillover components and short-term plasticity^3^. The tonic GABA_A_ receptor-mediated inhibitory conductance present in GC was also included^28^. This level of description reproduces the measured GC input-output relationship^29,30^. The synaptic connectivity of the experimentally constrained 3D network model was identical to the analytical model. A downstream decoder was trained to classify MF, GC, or shuffled GC spike counts in a 30 ms time window, corresponding to the effective integration time of GCs^3,30^. Despite the stochastic noise introduced by the Poisson input trains, networks with the sparse level of synaptic connectivity found in the cerebellar input layer separated MF activation patterns and sped up learning by up to 4-fold. Biologically detailed models also exhibited the same general trends for pattern separation and learning that were present in the analytical model: learning was fastest for sparsely connected networks, while densely connected networks performed worse than MFs (Fig. 6b). Moreover, the robustness of the normalized learning speed increased with increasing input correlations for sparsely connected networks, but did not significantly increase for densely connected networks.

**Figure 6.**
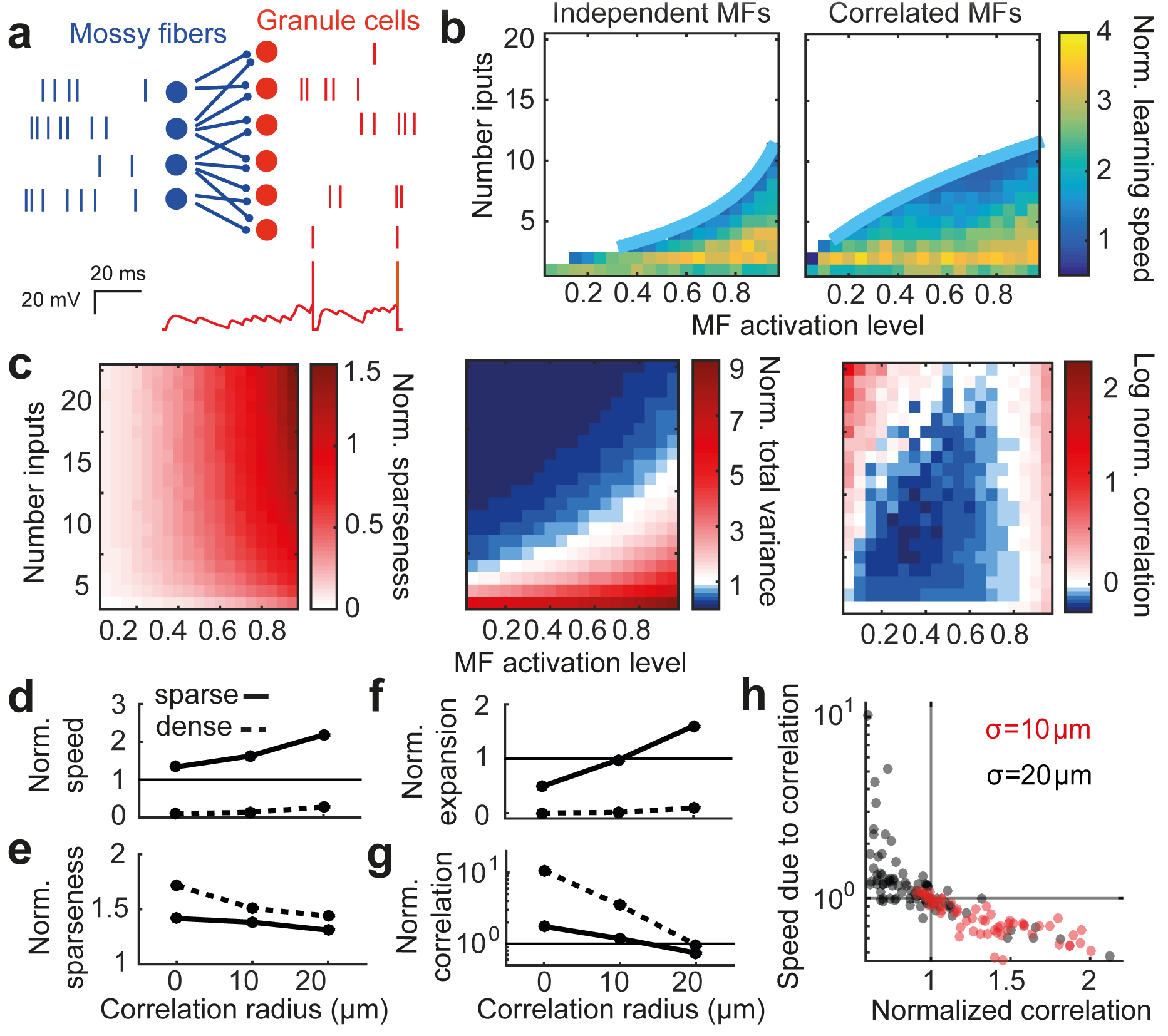
Pattern separation and learning speed depend on synaptic connectivity in biologically detailed spiking models of the cerebellar input layer. (**a**) Top: Schematic of biologically detailed spiking network model with sample spike trains. Bottom: example voltage trace from a granule cell (GC) in network. (**b**) Normalized learning speed for a spiking networks with independent (left) and correlated (right, σ = 20 μm) mossy fiber (MF) activation patterns. (**c**) Normalized sparseness (left), normalized total variance (middle), and log normalized population correlation (right) for networks with different numbers of synaptic connections receiving correlated MF activation patterns (σ = 20 μm). (d) Median normalized learning speed plotted against correlation radius for sparse (solid, N_syn_ = 4) and dense (dashed, N_syn_ = 16) synaptic connectivities. (e-g) Same as (**d**) for normalized sparseness (**e**), normalized total variance (**f**), and normalized population correlation (**g**). (**h**) Change in learning speed due to correlations (i.e., GC speed/ speed for shuffled GC spike trains) plotted against the normalized population correlation. Each point represents different values of N_syn_ and MF activation. Correlation radii were σ = 10 μm (red) or 20 μm (black).

To examine how sparsening of population activity and expansion of coding space contributed to the speed up in learning in biologically detailed models we first examined the normalized sparseness of the spike count patterns. The increase in the normalized sparseness with the number of synaptic inputs was more pronounced than for the simplified threshold-linear model (Fig. 6c, left). This is likely to be due to the dependence of the shape of the GC input-output function on the number of synaptic connections^3^. However, while the normalized learning speed increased with input correlations in sparsely connected networks (Fig. 6d), the normalized sparseness decreased (Fig. 6e). Therefore, sparseness was not able to explain the dependence of learning on the MF input correlation. In contrast, the normalized total variance had a similar dependence on synaptic connectivity and MF activation as the normalized learning speed (Fig. 6c, middle). Moreover, sparsely connected networks improved performance with increasing correlations while densely connected networks exhibited little change (Fig. 6F). However, the normalized total variance was unable to capture the full magnitude of the speedup for sparsely connected networks. Interestingly, decorrelation was more robust in the spiking models than in the corresponding reduced model (Fig. 6c, right). Like sparseness, this is likely to be due to the change in the GC input-output function with increasing numbers of inputs. In line with predictions from the analytical models, as the correlation radius increased, the normalized population correlation decreased (Fig. 6g). Finally, upon shuffling GC spike count patterns, we found a strong negative relationship between the population correlation and its impact on learning, with decorrelation speeding learning beyond the effects of expansion, as predicted by our analytical model (Fig. 6h c.f. Fig. 5e). These results show that the network connectivity and biophysical mechanisms present in the cerebellar input layer can implement effective pattern separation. Moreover they confirm the predictions from our simplified models, which show that sparse connectivity and nonlinear thresholding is essential for effective pattern separation and decorrelation in feedforward excitatory networks.

## Discussion

We have explored the relationship between the structure of excitatory feedforward networks and their ability to perform pattern separation. To do this we examined how both simplified and biologically detailed networks models with varying synaptic connectivity transform spatially correlated neural activity patterns, and how such a transformation affects the learning speed of a downstream classifier. Our results reveal that the structure of divergent feedforward networks governs pattern separation performance because increasing synaptic connectivity increases correlations in the output layer, counteracting the beneficial effects of expansion of coding space. Effective pattern separation is restricted to networks with few synaptic connections per neuron, since only sparsely connected networks are able to limit correlations introduced by the network and actively decorrelate input patterns through spike thresholding. Our work suggests that the evolutionarily conserved synaptic connectivity found in the input layer of the vertebrate cerebellum is optimal for separating spatially correlated input patterns, enabling faster learning in downstream circuits.

The idea that divergent feedforward networks separate overlapping activity patterns by expanding them into a higher dimensional space has a long history. In the cerebellum, pioneering work by Marr and Albus linked the structure of the input layer to expansion recoding of activity patterns^1,17^. Subsequent theoretical work has broadened our understanding of how pattern separation, information transfer, and learning can arise in cerebellar-like feedforward networks^2,3,6,31–33^. Our work extends these findings by quantifying how the synaptic connectivity within feedforward networks affects their ability to separate spatially correlated input patterns. We gained new insight into pattern separation by analyzing how network connectivity affects the size of the coding space, population sparseness and correlations, which are its key determinants. Moreover, we showed that biologically detailed network models with the sparse network connectivity and biophysical mechanisms present in the cerebellar input layer can decorrelate spatially correlated synaptic inputs, perform pattern separation, and speed up learning by a downstream classifier.

In addition to expansion recoding, classic studies have also highlighted the importance of sparse coding for pattern separation^1,4,17,31,33^. Moreover, our previous work showed that the sparse synaptic connectivity in the cerebellar input layer is well suited for performing lossless sparse encoding^3^. Our current findings provide new insight into pattern separation by showing that the increase in population sparseness from MFs to GCs is not able to explain the dependence of learning speed on MF correlation (Fig. 2a,b). While sparsening activity aids pattern separation for uncorrelated input patterns^4,6,31,33^, expansion of coding space and decorrelation of activity patterns had a greater impact on learning speed for more biologically realistic spatially correlated input patterns (Fig. 6c). Receptive field properties that are matched to the statistics of the sensory input could make expansion and sparse coding more effective by reducing the variance of sensory evoked activity patterns, thereby reducing pattern overlap^6^. But, excessive sparsening with high thresholds quenches the coding space (Fig. 4c) and results in a loss of information^3^, which are detrimental for robust pattern separation. Thus, for the random spatially correlated inputs studied here, expansion of coding space and decorrelation, rather than sparsening, are the main determinants of pattern separation in sparsely connected feedforward networks.

Inhibition has been shown to sparsen and decorrelate neural activity patterns^10,3,34,35^. Inhibition in the cerebellar input layer is relatively simple with a large fixed tonic GABA_A_ receptor-mediated inhibition of GCs (which was implemented in our biologically detailed simulations) that is complemented by a weaker activity-dependent component mediated by phasic release and GABA spillover from Golgi cells^28,36^. When network-activity dependent thresholding was included to approximate feedforward Golgi cell inhibition of GCs^3,37^, we observed greater decorrelation, but the dependence of pattern separation on network connectivity was preserved (Supplementary Fig. 2). Increasing the level of inhibition with the excitatory drive improves decorrelation because the higher spike threshold ensures that a substantial proportion of the correlated input remains subthreshold and is therefore filtered out. Dynamic inhibition is even more crucial for decorrelation in recurrent networks than simple feedforward networks, since the synaptic connectivity supports a finer balance of excitatory and inhibitory conductances, enabling more effective canceling out of inputs and stable asynchronous dynamics^38–41^.

Because pattern separation is essential for a wide range of sensory and motor processing, it is not surprising that divergent feedforward excitatory networks are found throughout the brain of both vertebrates and invertebrates. Interestingly, the connectivity in many of these networks is sparse. In the vertebrate auditory system, GCs in the dorsal cochlear nucleus receive only 3 synaptic connections^15^. GCs have 2-5 synapses in the input layer of the cerebellar-like electrosensory lateral line lobe of electric fish^42^. In the insect mushroom body, the numerous Kenyon cells receive olfactory information from an average of 7 random projection neurons^16^.

Furthermore, the characteristic 2-7 synaptic connections found in the cerebellar input layer has been evolutionarily conserved since the appearance of fish^43^. Our results indicate that such sparse connectivity is optimized for decorrelation and pattern separation and that this does not depend on the precise expansion ratio employed (Supplementary Fig. 1). These results are in agreement with recent analytical modeling, which predicts that the levels of sparse connectivity observed for cerebellar GCs (and Kenyon cells) are optimal for learning associations (A. Litwin-Kumar and K. Harris, personal communication). This suggests that the advantage of improved pattern separation and learning that sparse synaptic connectivity confers has been sufficient to conserve the structure of the cerebellar input layer for 400 million years.

MFs arise from multiple precerebellar nuclei in the brain stem and often project to specific regions in the cerebellar cortex, resulting in a large scale modular structure with regional specializations^44^. Within an individual module the receptive fields of MFs form a fractured map^45^, which at the local level, is likely to lead to spatially correlated activation. While single GC recordings suggest multimodal integration, in forelimb regions synaptic inputs can convey highly related information^46,47^, suggesting that the activity of MFs projecting to a local region are spatially correlated. Our results show that sparsely connected local networks with a 1:2.9 rosette to GC expansion ratio can decorrelate these input patterns enabling faster learning by downstream networks. Although only 1-2 rosettes are formed by a MF as it traverses a local network of 80 μm in diameter^48^, a complete MF axon forms 20-30 rosettes as it passes through an entire lobule^14^. At the level of a lobule the MF-GC expansion ratio is therefore substantially larger. Thus the improvement in pattern separation and learning conferred by the GC layer across an entire lobule could be significantly larger than the local network (e.g., see Supplementary Fig. 1 for a 1:9 expansion ratio).

Because MF inputs encode both discrete and continuous sensory variables, some areas of the cerebellum, such as the whisker system in Crus I/II, are likely to experience bouts of intense high frequency MF excitatory drive (100 – 1000 Hz), interspersed by quiescence^25,49^, while other areas such as vestibular and limb areas are likely to experience more slowly modulated sustained rate coded input at 10 -100 Hz^26,27^. Our findings suggest that the same network structure can perform efficient decorrelation and pattern separation for a wide range of MF excitatory drive. While there are likely to be region-specific specializations in synaptic properties and inhibition, the uniformity of input layer structure suggest that it acts as a generic preprocessing unit that decorrelates and separates dense MF activation patterns, enabling faster associative learning in the molecular layer.

A core function of the cerebellar cortex is to learn the sensory consequences of motor actions, allowing it to predict and refine motor action and to enable animals to perform sensory processing in the context of active movement^50–52
^. In Purkinje cells, which receive ~200,000 GC inputs, learning is achieved by altering synaptic strength via LTD and LTP, depending on when parallel fiber activity occurs in relation to the climbing fiber input^53^. We used perceptron-based learning to assay pattern separation and learning performance, since early work recognized analogies between supervised learning in Purkinje cells and perceptrons^17^. More recent work has shown that perceptron learning is consistent with LTD at the parallel fiber-Purkinje cell synapse^54^ and that the timing of learning rules can be tuned to account for feedback delays in the PF input^55^. While these factors support our use of perceptron-based learning as an assay of learning performance, important functional differences with Purkinje cells limit finer-grained insights into cerebellar learning using a perceptron-based approach. These include the fact that Purkinje cells fire spontaneously at 40-90 Hz, receive strong phasic inhibitory inputs and exhibit mGluR7 mediated modulation of intrinsic potassium conductances which introduce pauses in their firing rate^56,57^. Nevertheless, decorrelation of GC activity is expected to improve the performance of the conversion of spatial activity into temporal sequences via these mechanisms.

While encoding temporal sequences is clearly essential to motor control it is not clear whether the plasticity mechanisms that underlie such function fit better within an extended classical framework of associative learning^33^, temporal encoding^58^, or implementing an adaptive filter^59^. Irrespective of which of these models the cerebellar cortex implement, our results highlight the detrimental effect that spatial correlations can have on sensorimotor learning and the substantial improvements in performance that can be achieved through spatial decorrelation and expansion of coding space. Moreover, our results show that the input layer performs effective pattern separation across a wide range of spatial correlations and input activity and does not require the exceedingly sparse coding regimes envisioned by Marr^1^. Indeed, the input layer could improve learning when up to 66% of GCs were active and could decorrelate inputs when up to 33% of GCs were active. While our study focused on the importance of spatial correlations, it will be interesting to investigate whether comparable improvements can be obtained in the temporal domain, since temporal expansion and sparsening will increase the dimensionality of the system further^9,60–62^.

Our results are consistent with several existing experimental manipulations in the cerebellar input layer. Reduction in the number of functional GCs by 90% using a genetic manipulation that blocked their output resulted in deficits in the consolidation of motor learning^63^. Our findings suggest that this phenotype arose from deficits in pattern separation and learning speed due to the reduced coding space available, although concomitant changes in long-term plasticity could also contribute. Another prediction is that decreasing the threshold of GCs will affect the expansion-correlation tradeoff and reduce pattern separation and impair learning. Interestingly, lowering the spike threshold by specifically deleting the KCC2 chloride transporter in cerebellar GCs, resulted in impaired learning consolidation^64^. Similarly, inhibiting a negative feedback circuit in the drosophila olfactory system increased correlations in odor-evoked activity patterns and impaired odor discrimination^10^. Both these findings are consistent with our prediction that lowering threshold or removing feedforward inhibition will increase correlations in feedforward networks and impair pattern separation and learning.

The most direct experimental tests of the predictions of this work is that spiking in GCs should have a larger total variance and smaller population correlation than spiking in MF inputs. However, our results show that network decorrelation cannot be revealed by measuring pairwise correlations, consistent with previous work that showed pairwise measurements can underestimate collective activity in larger populations^65^. Our analysis indicates that dense recordings from a large fraction of the neurons in the local network are required to measure population correlation in MFs and GCs (Fig. 3e). Recent developments in high speed random access 3D two-photon imaging^66,67^ and sensitive genetically encoded Ca^2+^ indicators ^68^ potentially make this type of challenging measurement feasible for the first time. Application of these new technologies would provide direct experimental tests of our findings, thereby improving our understanding of how spatially correlated activity patterns are transformed and separated in the cerebellar cortex.

## Methods

**Anatomical network model**. Both the analytical and biophysical models used an experimentally constrained anatomically realistic network connectivity model of an 80 μm diameter ball of within the granular layer^3^. MF rosettes and GCs were positioned according to their observed densities. GCs were connected to a fixed number (N_syn_) of MFs, which were chosen randomly while constraining the MF-GC distance to be as close as possible to 15 μm, the average dendritic length.

**Spatially correlated input patterns**. MF activation patterns were created using a method based on Dichotomized Gaussian models that generates binary vectors with specified mean firing rates (MF activation levels) and correlations^20^. The correlation coefficient between two MF patterns was chosen to be a Gaussian function of distance with the correlation radius parameterized by its standard deviation σ. For the analytical model, these binary patterns were used directly. For the detailed model, activated MFs fired at 50 Hz while inactivated MFs were silent.

**Simplified network model.** GC activity was given by:

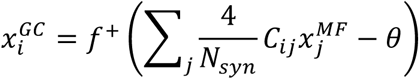

Where N_syn_ is the number of synaptic inputs per GC, C_ij_ is the binary connectivity matrix determined by the anatomical network model, and *f*^+^ is a rectified-linear function, i.e., *f*^+^(x) = *max*(0, x). Unless otherwise specified, the threshold was set to θ = 3, in line with experimental evidence that three MFs on average are required to generate a spike in GC^29,69^.

**Biologically detailed network model.** MFs were modeled as modified Poisson processes with a 2 ms refractory period and firing rate determined by the generated binary activity patterns described above (50 Hz if the MF was activated, silent otherwise). GCs were based on a previously published model of integrate-and-fire neurons with experimentally measured passive properties and experimentally constrained AMPA and NMDA conductances, short-term plasticity and spillover components as well as constant GABA conductance representing tonic inhibition^3^ (see Supplementary Methods). The model was written in NeuroML2 and simulated in jLEMS^70^. For learning and population-level analysis, activity patterns were defined as the vector of spike counts in a 30 ms window (after discarding an initial 150 ms period to reach steady state).

**Implementation of perceptron learning.** A perceptron decoder was trained to classify 640 input patterns into 10 random classes. Random classification was chosen to ensure maximal overlap between patterns. The number of classes was chosen to be slightly under the memory capacity for a wide range of parameters, allowing comparison of learning in different networks for a relatively complex task. Online learning was implemented with backpropagation learning on a single layer neural network with sigmoidal nodes and a small fixed learning rate of 0.01. The inputs consisted of either the raw MF or the GC activity patterns. Learning took place over 5000 epochs, each of which consisted of presentations of all 640 patterns in a random order. Learning speed was defined as 1/N_E_, where N_E_ is the number of training epochs until the root-mean-square error reached a threshold of 0.2. Other error thresholds gave qualitatively similar results.

**Analysis of activity patterns.** Population sparseness was measured as^21^:

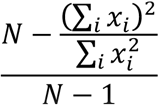

Where N is the number of neurons and x_i_ is the ith neuron’s activity (simplified model) or spike count (detailed model). The above quantity was averaged over all activity patterns. To quantify pattern expansion, we use the total variance, i.e. the sum of all variances:

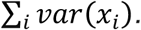

We defined the population correlation as:

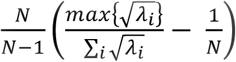

Where λ_*i*_ are the eigenvalues of the covariance matrix. The first term in this expression describes how elongated the distribution is in its principal direction. The second term subtracts the value 1/N so that an uncorrelated homogenous Gaussian would have a value of zero. A modified version of the population correlation to control for heterogeneous variances did not affect the results (see Supplementary Methods). Finally, the scaling factor of 
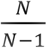
 normalizes the expression so that its maximum value is 1.

**Partial shuffling of spiking activity.** We developed a shuffling technique to increase or decrease the population correlation to a desired level, while keeping the mean and variance of each neuron fixed. To shuffle patterns to a lower level of correlation, for each neuron in the population we swapped the spike counts for random pairs of patterns. This was iterated over the full population and over random pattern pairs until the resulting activity patterns had the desired population correlation. Conversely, to shuffle activity patterns in a way that would increase correlations, we took random pairs of patterns and swapped the activity so that each cell had a lower spike count for the first pattern and higher activity for the second pattern. This procedure modifies the activity patterns so that the population overall tends to be more active together. We then tested perceptron learning based on the new shuffled activity patterns. See Supplementary Methods for additional details.

**Code availability.** Models and scripts for running simulations will be made available at https://github.com/SilverLabUCL/ upon publication.

## Acknowledgements

This work was supported by the Wellcome Trust (095667; 203048; 101445). RAS in receipt of a Wellcome Trust Principal Research Fellowship and an ERC advanced grant (294667). We thank Eugenio Piasini, Sadra Sadeh, Yann Sweeney, Antione Valera and Tommy Younts for comments on the manuscript and Ashok Litwin-Kumar and Kam Harris for helpful discussions.

## Author contributions

N. A. C. G. carried out the simulations and analyzed the data. N. A. C. G., C. C., and R. A. S. conceived the project and designed the experiments. N. A. C. G and R. A. S wrote the manuscript.

## Competing financial interests

The authors declare no competing financial interests.

## References

1. Marr, D. A theory of cerebellar cortex. J. Physiol. 202, 437–470 (1969).

2. Kanerva, P. in Associative Neural Memories: Theory and Implementation 50–76 (1993).

3. Billings, G., Piasini, E., Lorincz, A., Nusser, Z. & Silver, R. A. Network Structure within the Cerebellar Input Layer Enables Lossless Sparse Encoding. Neuron 83, 960–974 (2014).

4. Olshausen, B. A. & Field, D. J. Emergence of simple-cell receptive field properties by learning a sparse code for natural images. Nature 381, 607–609 (1996).

5. Földiak, P. in The Handbook of Brain Theory and Neural Networks (ed. Arbib, M. A.) 1064–1068 (MIT Press, 2002). doi:10.1.1.16.1720.

6. Babadi, B. & Sompolinsky, H. Sparseness and Expansion in Sensory Representations. Neuron 83, 1213–1226 (2014).

7. Friedrich, R. W. Neuronal computations in the olfactory system of zebrafish. Annu. Rev. Neurosci. 36, 383–402 (2013).

8. Gschwend, O. et al. Neuronal pattern separation in the olfactory bulb improves odor discrimination learning. Nat. Neurosci. 18, 1474–1482 (2015).

9. Laurent, G. Olfactory network dynamics and the coding of multidimensional signals. Nat. Rev. Neurosci. 11, 884–895 (2002).

10. Lin, A. C., Bygrave, A. M., de Calignon, A., Lee, T. & Miesenböck, G. Sparse, decorrelated odor coding in the mushroom body enhances learned odor discrimination. Nat. Neurosci. 17, 559–68 (2014).

11. Oertel, D. & Young, E. D. What’s a cerebellar circuit doing in the auditory system? Trends in Neurosciences 27, 104–110 (2004).

12. Yassa, M. A. & Stark, C. E. L. Pattern separation in the hippocampus. Trends in Neurosciences 34, 515–525 (2011).

13. Leutgeb, J. K., Leutgeb, S., Moser, M.-B. & Moser, E. I. Pattern Separation in the Dentate Gyrus and CA3 of the Hippocampus. Science (80-.). 315, 961–966 (2007).

14. Eccles, J. C., Ito, M. & Szentágothai, J. The cerebellum as a neuronal machine. (Springer, 1967).

15. Mugnaini, E., Osen, K. K., Dahl, A. L., Friedrich, V. L. & Korte, G. Fine structure of granule cells and related interneurons (termed Golgi cells) in the cochlear nuclear complex of cat, rat and mouse. J. Neurocytol. 9, 537–570 (1980).

16. Caron, S. J. C., Ruta, V., Abbott, L. F. & Axel, R. Random convergence of olfactory inputs in the Drosophila mushroom body. Nature 497, 113–117 (2013).

17. Albus, J. S. A Theory of Cerebellar Function. Math. Biosci. 10, 25–61 (1971).

18. Baker, J. L. Is there a support vector machine hiding in the dentate gyrus? Neurocomputing 52–54, 199–207 (2003).

19. Ganguli, S. & Sompolinsky, H. Statistical Mechanics of Compressed Sensing. Phys. Rev. Lett. 104, 188701 (2010).

20. Macke, J. H., Berens, P., Ecker, A. S., Tolias, A. S. & Bethge, M. Generating spike trains with specified correlation coefficients. Neural Comput. 21, 397–423 (2009).

21. Vinje, W. E. & Gallant, J. L. Sparse Coding and Decorrelation in Primary Visual Cortex During Natural Vision. Science (80-.). 287, 1273–1276 (2000).

22. de la Rocha, J., Doiron, B., Shea-Brown, E., Josić, K. & Reyes, A. D. Correlation between neural spike trains increases with firing rate. Nature 448, 802–806 (2007).

23. Wiechert, M. T., Judkewitz, B., Riecke, H. & Friedrich, R. W. Mechanisms of pattern decorrelation by recurrent neuronal circuits. Nat. Neurosci. 13, 1003–1010 (2010).

24. Nirenberg, S. & Latham, P. E. Population coding in the retina. Current Opinion in Neurobiology 8, 488–493 (1998).

25. Rancz, E. A. et al. High-fidelity transmission of sensory information by single cerebellar mossy fibre boutons. Nature 450, 1245–1248 (2007).

26. Arenz, A., Silver, R. A., Schaefer, A. T. & Margrie, T. W. The contribution of single synapses to sensory representation in vivo. Science (80-.). 321, 977–980 (2008).

27. Van Kan, P. L., Gibson, A. R. & Houk, J. C. Movement-related inputs to intermediate cerebellum of the monkey. J. Neurophysiol. 69, 74–94 (1993).

28. Farrant, M. & Nusser, Z. Variations on an inhibitory theme: phasic and tonic activation of GABA(A) receptors. Nat. Rev. Neurosci. 6, 215–229 (2005).

29. Schwartz, E. J. et al. NMDA receptors with incomplete Mg2+ block enable low-frequency transmission through the cerebellar cortex. J. Neurosci. 32, 6878–6893 (2012).

30. Rothman, J. S., Cathala, L., Steuber, V. & Silver, R. A. Synaptic depression enables neuronal gain control. Nature 457, 1015–1018 (2009).

31. Tyrrell, T. & Willshaw, D. Cerebellar cortex: its simulation and the relevance of Marr’s theory. Philos. Trans. R. Soc. Lond. B. Biol. Sci. 336, 239–257 (1992).

32. Torioka, T. Pattern Separability and the Effect of the Number of Connections in a Random Neural Net with Inhibitory Connections. Biol. Cybern. 31, 27–35 (1978).

33. Schweighofer, N., Doya, K. & Lay, F. Unsupervised learning of granule cell sparse codes enhances cerebellar adaptive control. Neuroscience 103, 35–50 (2001).

34. Tetzlaff, T., Helias, M., Einevoll, G. T. & Diesmann, M. Decorrelation of Neural-Network Activity by Inhibitory Feedback. PLoS Comput. Biol. 8, (2012).

35. King, P. D., Zylberberg, J. & DeWeese, M. R. Inhibitory interneurons decorrelate excitatory cells to drive sparse code formation in a spiking model of V1. J. Neurosci. 33, 5475–5485 (2013).

36. Duguid, I., Branco, T., London, M., Chadderton, P. & Hausser, M. Tonic Inhibition Enhances Fidelity of Sensory Information Transmission in the Cerebellar Cortex. J. Neurosci. 32, 11132–11143 (2012).

37. Kanichay, R. T. & Silver, R. A. Synaptic and cellular properties of the feedforward inhibitory circuit within the input layer of the cerebellar cortex. J. Neurosci. 28, 8955–8967 (2008).

38. Okun, M. & Lampl, I. Instantaneous correlation of excitation and inhibition during ongoing and sensory-evoked activities. Nat. Neurosci. 11, 535–537 (2008).

39. van Vreeswijk, C. & Sompolinsky, H. Chaos in neuronal networks with balanced excitatory and inhibitory activity. Science (80-.). 274, 1724–1726 (1996).

40. Renart, A. et al. The asynchronous state in cortical circuits. Science (80-.). 327, 587–90 (2010).

41. Graupner, M. & Reyes, A. D. Synaptic Input Correlations Leading to Membrane Potential Decorrelation of Spontaneous Activity in Cortex. J. Neurosci. 33, 15075–15085 (2013).

42. Kennedy, A. et al. A temporal basis for predicting the sensory consequences of motor commands in an electric fish. Nat. Neurosci. 17, 416–422 (2014).

43. Wittenberg, G. M. & Wang, S. S. H. in Dendrites (eds. Stuart, G., Spruston, N. & Häusser, M.) 43–67 (Oxford University Press, 2007). doi:10.1093/acprof:oso/9780198566564.003.0002

44. Apps, R. & Hawkes, R. Cerebellar cortical organization: a one-map hypothesis. Nat. Rev. Neurosci. 10, 670–681 (2009).

45. Shambes, G. M., Gibson, J. M. & Welker, W. Fractured somatotopy in granule cell tactile areas of rat cerebellar hemispheres revealed by micromapping. Brain. Behav. Evol. 15, 94–140 (1978).

46. Bengtsson, F. & Jörntell, H. Sensory transmission in cerebellar granule cells relies on similarly coded mossy fiber inputs. Proc. Natl. Acad. Sci. U. S. A. 106, 2389–2394 (2009).

47. Powell, K., Mathy, A., Duguid, I. & Häusser, M. Synaptic representation of locomotion in single cerebellar granule cells. Elife 4, e07290 (2015).

48. Sultan, F. Distribution of mossy fibre rosettes in the cerebellum of cat and mice: Evidence for a parasagittal organization at the single fibre level. Eur. J. Neurosci. 13, 2123–2130 (2001).

49. Ritzau-Jost, A. et al. Ultrafast action potentials mediate kilohertz signaling at a central synapse. Neuron 84, 152–163 (2014).

50. Proville, R. D. et al. Cerebellum involvement in cortical sensorimotor circuits for the control of voluntary movements. Nat. Neurosci. 17, 1233–1239 (2014).

51. Brooks, J. X., Carriot, J. & Cullen, K. E. Learning to expect the unexpected: rapid updating in primate cerebellum during voluntary self-motion. Nat. Neurosci. 18, 1310–1317 (2015).

52. Wolpert, D. M., Miall, R. C. & Kawato, M. Internal models in the cerebellum. Trends Cogn. Sci. 2, 338–347 (1998).

53. Gao, Z., Beugen, B. J. & Van De Zeeuw, C. I. Distributed synergistic plasticity and cerebellar learning. Nat. Rev. Neurosci. 13, 619–635 (2012).

54. Brunel, N., Hakim, V., Isope, P., Nadal, J.-P. & Barbour, B. Optimal information storage and the distribution of synaptic weights: Perceptron versus Purkinje cell. Neuron 43, 745–757 (2004).

55. Suvrathan, A., Payne, H. L. & Raymond, J. L. Timing Rules for Synaptic Plasticity Matched to Behavioral Function. Neuron 92, 959–967 (2016).

56. Steuber, V. et al. Cerebellar LTD and Pattern Recognition by Purkinje Cells. Neuron 54, 121–136 (2007).

57. Johansson, F., Carlsson, H. A. E., Rasmussen, A., Yeo, C. H. & Hesslow, G. Activation of a Temporal Memory in Purkinje Cells by the mGluR7 Receptor. Cell Rep. 13, 1741–1746 (2015).

58. De Zeeuw, C. I. et al. Spatiotemporal firing patterns in the cerebellum. Nat. Rev. Neurosci. 12, 327–344 (2011).

59. Dean, P., Porrill, J., Ekerot, C.-F. & Jörntell, H. The cerebellar microcircuit as an adaptive filter: experimental and computational evidence. Nat. Rev. Neurosci. 11, 30–43 (2010).

60. Chabrol, F. P., Arenz, A., Wiechert, M. T., Margrie, T. W. & DiGregorio, D. A. Synaptic diversity enables temporal coding of coincident multisensory inputs in single neurons. Nat. Neurosci. 18, 718–727 (2015).

61. Medina, J. F. & Mauk, M. D. Computer simulation of cerebellar information processing. Nat. Neurosci. 3, 1205–1211 (2000).

62. Buonomano, D. V & Maass, W. State-dependent computations: spatiotemporal processing in cortical networks. Nat. Rev. Neurosci. 10, 113–125 (2009).

63. Galliano, E. et al. Silencing the Majority of Cerebellar Granule Cells Uncovers Their Essential Role in Motor Learning and Consolidation. Cell Rep. 3, 1239–1251 (2013).

64. Seja, P. et al. Raising cytosolic Cl-in cerebellar granule cells affects their excitability and vestibulo-ocular learning. EMBO J. 31, 1217–30 (2012).

65. Schneidman, E., Berry, M. J., Segev, R. & Bialek, W. Weak pairwise correlations imply strongly correlated network states in a neural population. Nature 440, 1007–1012 (2006).

66. Nadella, K. M. N. S. et al. Random access scanning microscopy for 3D imaging in awake behaving animals. Nat. Methods 13, 1001–1004 (2016).

67. Ji, N., Freeman, J. & Smith, S. L. Technologies for imaging neural activity in large volumes. Nat. Neurosci. 19, 1154–64 (2016).

68. Chen, T.-W. et al. Ultrasensitive fluorescent proteins for imaging neuronal activity. Nature 499, 295–300 (2013).

69. Jörntell, H. & Ekerot, C.-F. F. Properties of somatosensory synaptic integration in cerebellar granule cells in vivo. J. Neurosci. 26, 11786–11797 (2006).

70. Cannon, R. C. et al. LEMS: a language for expressing complex biological models in concise and hierarchical form and its use in underpinning NeuroML 2. Front. Neuroinform. 8, (2014).

